# GCI: a continuity inspector for complete genome assembly

**DOI:** 10.1101/2024.04.06.588431

**Authors:** Quanyu Chen, Chentao Yang, Guojie Zhang, Dongya Wu

## Abstract

**Motivation:** Recent advances in long-read sequencing technologies have significantly facilitated the production of high-quality genome assembly. The telomere-to-telomere (T2T) gapless assembly has become the new golden standard of genome assembly efforts. Several recent efforts have claimed to produce T2T level reference genomes. However, a universal standard is still missing to qualify a genome assembly to be at T2T standard. Traditional genome assembly assessment metrics (N50 and its derivatives) have no capacity in differentiate between nearly T2T assembly and the truly T2T assembly in continuity either globally and locally. Also these metrics are independent of raw reads, which make them inflated easily by artificial operations. Therefore a gaplessness evaluation tool at single nucleotide resolution to reflect true completeness is urgently needed in the era of complete genomes.

**Results:** Here, we present a tool called Genome Continuity Inspector (GCI) to assess genome assembly continuity at the single base resolution, that can evaluate how close a genome assembly is close to T2T level. GCI utilized multiple aligners to map long reads from multiple platforms back to the assembly. By incorporating curated mapping coverage of high-confidence read alignments, GCI identifies potential assembly issues. Meanwhile, it also reports GCI scores to quantify the assembly overall continuity in the whole genome or chromosome scale.

**Availability and implementation:** The open-source GCI code is freely available on Github (https://github.com/yeeus/GCI) under the MIT license.

## Introduction

Long-read sequencing technologies based on Pacbio HiFi and Oxford Nanopore Technology (ONT) sequencing platform have been routinely used in *de novo* genome assembling pipelines. It has been demonstrated in several gapless genome assemblies, including human (Nurk et al., 2023; Yang et al., 2023), chicken (Huang et al., 2023), *Arabidopsis thaliana* (Naish et al., 2021), and rice (Song et al., 2022) that previous assembly challenge on highly repetitive regions can now be largely filled using long reads. A series of metrics are currently used to evaluate the quality of *de novo* genome assemblies based on “3C criterion” (completeness, correctness and continuity)(Wang P. & Wang F., 2023). For completeness, BUSCO, CEGMA and similar gene-mapping based tools (e.g. asmgene) are widely used (Li, 2018). These gene-focus assessments however can not represent the quality of gene-desert regions with complex structures. *K*-mer completeness evaluated by Merqury provides another completeness indicator, but is sensitive to low read quality and experimental contamination, which would introduce false or exogenous *K*-mers (Arang et al., 2020). For correctness, QV (consensus quality) is widely used to measure shared *K*-mers between raw reads and final assembly but can be artificially manipulated by removing erroneous assembly sequences (Arang et al., 2020).

Genome assembly continuity is typically measured by the metric contig N50, which represents the length of the shortest contig at half of total assembly sequence length. However, contig N50 and its derivative NG50, auN (https://lh3.github.io/2020/04/08/a-new-metric-on-assembly-contiguity) or E-size values (Salzberg et al., 2012) have easily reached or approaching their theoretical maximums due to the nature of contig N50’s discontinuity, suggesting limited capacity in differentiating among inter-individual assemblies using long reads or reflecting assembly improvement. In other words, once an assembly contig N50 has reached the value of chromosome N50 length, any improvement in gap filling can not be reflected by contig N50 any more. Furthermore, assembly continuity could be artificially inflated by replacing or removing gaps directly, which can not be detected solely from the assembly sequences. Therefore the tools to detect assembly errors at base-level resolution by realigning raw reads back have been urgently required to ensure the authenticity of a truly gapless assembly.

Mapping long reads back against genome assembly can reveal abnormal signals (e.g. mapping quality, clipping information, read coverage, edit distance/mismatch), which could be used to identify potential assembly errors. To detect base-resolution assembly errors, some tools have been developed based on this strategy, including Flagger (Liao et al., 2023) and CRAQ (Li et al., 2023). The T2T-polish pipeline developed by human genome T2T consortium (hereafter called as T2T-polish) also includes a sub-module with this function (Cartney et al., 2022).

Flagger was developed by Human Pangenome Reference Consortium (HPRC), which has been applied in the human pangenome study (Liao et al., 2023). It detects the anomalies in the read coverage and partitions the assembly into different categories predicting the accuracy of the assembly, such as duplicated, collapsed, erroneous blocks. T2T-polish pipeline also reports the assembly issue regions primarily based on read mapping coverage. CRAQ (Clipping information for Revealing Assembly Quality) uses clipping information of read alignments to detect potential assembly errors, but ignores the regions with extremely low or high coverage (Li et al., 2023). Briefly, abnormal mapping signals are collected as assembly issues based on reads coverage for Flagger and T2T-polish and clipping information for CRAQ. Such approach is understandable, but except assembly errors, reads sequencing bias in different genomic regions and aligning bias in highly repetitive regions by different aligners will also result in the occurrence of mapping anomalies, which would inevitably report a certain number of false positives in detecting assembly issues.

Here, we present a new alignment-based evaluator called Genome Continuity Inspector (GCI) for genome assembly assessment, particularly for evaluation of the assembly quality of genomes at T2T level or near T2T level. GCI integrates alignments of long reads generated from multiple platforms back to the assembly using multiple aligners. Unlike the strategy to detect issues using abnormal signals of reads mapping, it calls potential assembly issues based on curated coverage of high-confidence read alignments. Additionally, GCI calculates scores to quantify the overall continuity of a genome assembly at the genome or chromosome level. Briefly, GCI provides a new strategy to evaluate the quality of genome assembly, particularly in the T2T era.

## Materials and Methods

### Overview of GCI

#Reads mapping and filtering: GCI is a computational pipeline that uses alignment files (BAM or PAF format), generated by mapping long reads (HiFi and Nanopore reads) back to the final assembly as inputs, and outputs an score as an indicator of assembly continuity and reports the potential assembly issues. The tool requires alignments that pass stringent filtering criteria. All unmapped, secondary and supplementary alignments are discarded. Moreover, mapping quality (<30 in default), mapping identity (<90% in default) and clipped proportion (>10% in default) are further employed to remove low-quality alignments. Considering the potential alignment bias produced by mapping algorithms among aligners, at least two popular sequence aligners (e.g. minimap2 (Li, 2018), Winnowmap2 (Jain et al., 2022), VerityMap (Bzikadze et al., 2022)) are recommended to run on the same data (Figure 1). The alignments generated by the two aligners which pass the mapping quality requirements and have consistent mapping coordinates (overlap ≥90% in default) are kept. The mapping accuracy and sensitivity are prominently different between aligners. For example, minimap2 runs much faster than Winnowmap2 but underperforms in alignment of highly repetitive sequences, with low mapping quality (usually approaching to zero) (Jain et al., 2022). Therefore to rescue alignments in repetitive regions, a read is kept if one aligner produces a high mapping quality (≥50 in default) for this read.

**Figure 1.**
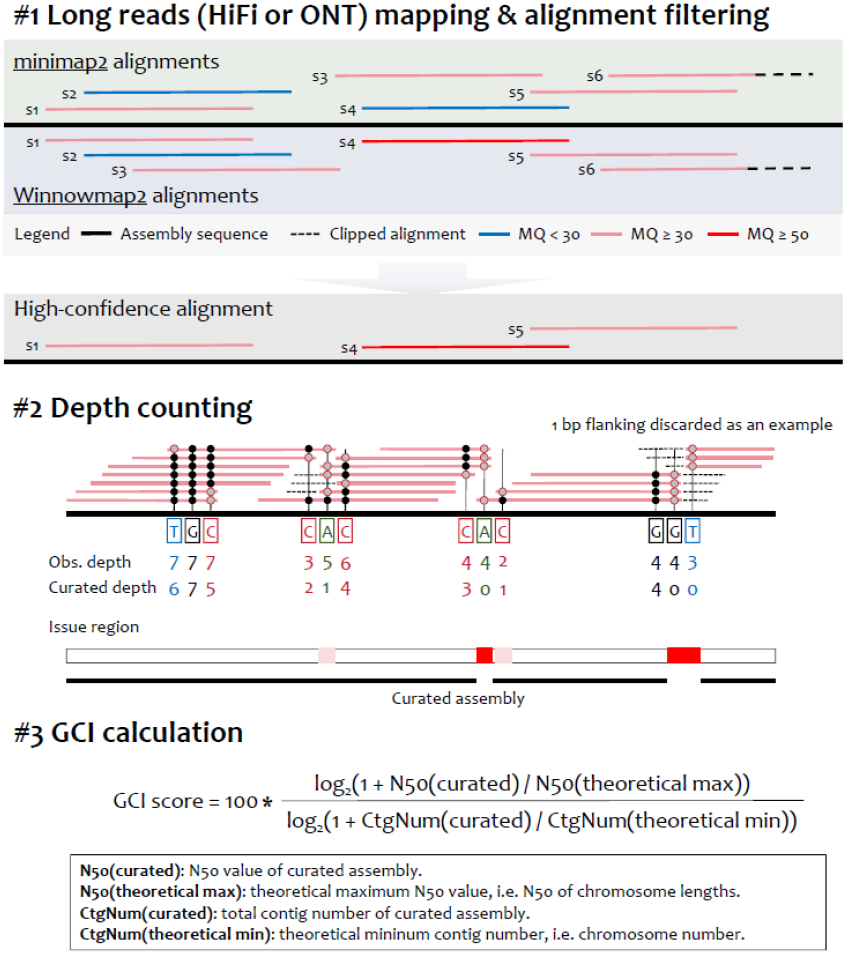
Workflow of GCI. Multiple aligning strategies (e.g. minimap2 and Winnowmap2) are used in mapping HiFi or ONT long reads against assembly sequence. Each read has two alignment results. After a series of stringent integrating and filtering, high-confidence alignments are curated and kept. By trimming both ends of the read alignment, curated depth is counted for each reference base. Potential assembly issues are then defined based on zero or extremely low depth. A curated assembly is produced by replacing assembly issues with gaps. To profile overall genome-wide continuity of assembly, GCI scores are calculated by considering both the contig N50 values and contig numbers of the curated assembly and theoretically gapless assembly. Obs., observed; CtgNum, contig number.

#Depth counting: After a series of strict alignment filtration, we count the mapping coverage for each base. Rather than directly counting mapping depth using samtools depth (observed depth), GCI workflow firstly trims the alignments of each flanking (one base as an example in Figure 1), to rule out clipped alignments and improve the depth estimation.

#GCI score calculating: According to the curated mapping depth for each base, GCI reports the potential assembly issues, where the regions have zero or extremely low depth. Physically close issues (e.g. distance less than 0.5% of chromosome length) are merged. The original chromosomes or sequences are subsequently split into curated contigs at loci with no read alignment supporting. Curated contig N50 and number are calculated for the curated assembly. Finally, considering the discontinuity of contig N50, GCI integrates both contig N50 value and contig number of curated assembly and quantifies the gap of assembly continuity to a truly gapless T2T assembly, using a GCI score (scaled from zero to 100) (Figure 1). Even if the contig N50 value has been saturated, the contig numbers could be used to quantify the continuity differences between assemblies. For a true T2T assembly, no issues or gaps would be detected and thus the curated contig N50 equals to the theoretical maximum N50 (chromosome N50) and the contig number equals to the number of chromosomes, which will thus produce a GCI score of 100 for such an assembly.

#Output: Potential assembly issue regions with zero or low-depth read alignment supports, and GCI scores for whole genome assembly and each chromosome are reported. Additionally, curated mapping depth plots are available for manual check.

### Benchmarking datasets

Several high-quality genomes have been released recently and some claimed to be at T2T level or near T2T, including several human genomes (CHM13 (Nurk et al., 2023), CN1 (Yang et al., 2023) and HG002 (Javis et al., 2023)), and other model organisms (chicken (GGswu (Huang et al., 2023)), *Arabidopsis* (Col-CEN, Naish et al., 2021) and rice (MH63)(Song et al., 2022)). To demonstrate the performance of GCI workflow in assessing quality of genome assembly, we downloaded the genome assemblies and corresponding raw long reads (HiFi and ONT) of these genomes and performed the assessment with GCI. All HiFi reads were firstly filtered using HiFiAdapter (Sim et al., 2022). For a haploid (i.e. CHM13), highly homozygous or self-fertilized (i.e. Col-CEN and MH63) or unphased (i.e. GGswu) diploid assembly, long reads were mapped against corresponding assembly directly. For haplotype-resolved assembly of the diploid genomes (i.e. human genomes CN1 and HG002), ONT and HiFi reads were firstly phased into paternal and maternal haplotypes based on parental genomic information using canu (Nurk, et al., 2020). The unphased reads of HiFi and ONT data were randomly and averagely assigned to the two haplotypes. Due to the lack of chromosome Y in CHM13, chromosome Y was not evaluated for all three human genomes. Plastid (mitochondria and chloroplast) genomes were also removed before aligning.

### Comparison among assembly issue detection tools

Two base-resolution quality evaluators, CRAQ (https://github.com/JiaoLaboratory/CRAQ) and T2T-polish (https://github.com/arangrhie/T2T-Polish) were used to detect potential assembly issues for the genomes used in this study and compared against GCI’s performance. For GCI, alignment BAM or PAF files generated by minimap2 and Winnowmap2 using all available long reads (HiFi and ONT) were input. CRAQ requires a single long read alignment as input, thus we provided the ONT read alignments produced by Winnowmap2, given the superior assembly continuity provided by ONT long reads. As recommended by CRAQ, NGS read alignments were also provided. For T2T-polish pipeline, HiFi and ONT read alignment files were input independently to run and the resulting issue regions were integrated to define a final dataset of assembly issues. Default parameters were used in both CRAQ and T2T-polish analyses.

## Results

### GCI score shows higher sensitivity in evaluating assembly continuity for high-quality genomes

We explored the sensitivity in assessing assembly continuity of GCI score, contig N50 and its derive auN from public released genomes and simulated data. To make them comparable, we firstly scaled contig N50 and auN values as scores from zero to 100 by their theoretical maximum values, respectively. CHM13 was the first human genome to be completely assembled, with multiple versions of updates. Most gaps (84/89) in v0.7 were filled in v0.9 and the remaining five rDNA gaps were resolved in v1.1 (https://github.com/marbl/CHM13). Contig N50 and auN had achieved their theoretical maximums since v0.9 and the continuity improvement by filling rDNA gaps was not reflected by them, while GCI score captured this change (Figure 2A). Additionally, we calculated these three metrics for population-level human phased genome assemblies from HGSVC (The Human Genome Structural Variation Consortium) phase 3. Given similar contig N50 scores, GCI scores showed greater deviations than those of auN, which suggesting GCI’s higher capacity in distinguishing continuity (Figure 2B). By randomly simulating gaps in the genome of CHM13, we found that contig N50 was the most insensitive and GCI score exhibited highest rate of decline as gap number increased, particularly for the nearly complete assemblies with limited gaps (Figure 2C). Moreover, higher slope was observed for GCI than auN when the contig N50 gradually approached to its maximum (Figure 2D). In other words, GCI score had a better ability to amplify and distinguish continuity for high-quality assemblies, in either reflecting assembly improvement or comparing inter-individual quality.

**Figure 2.**
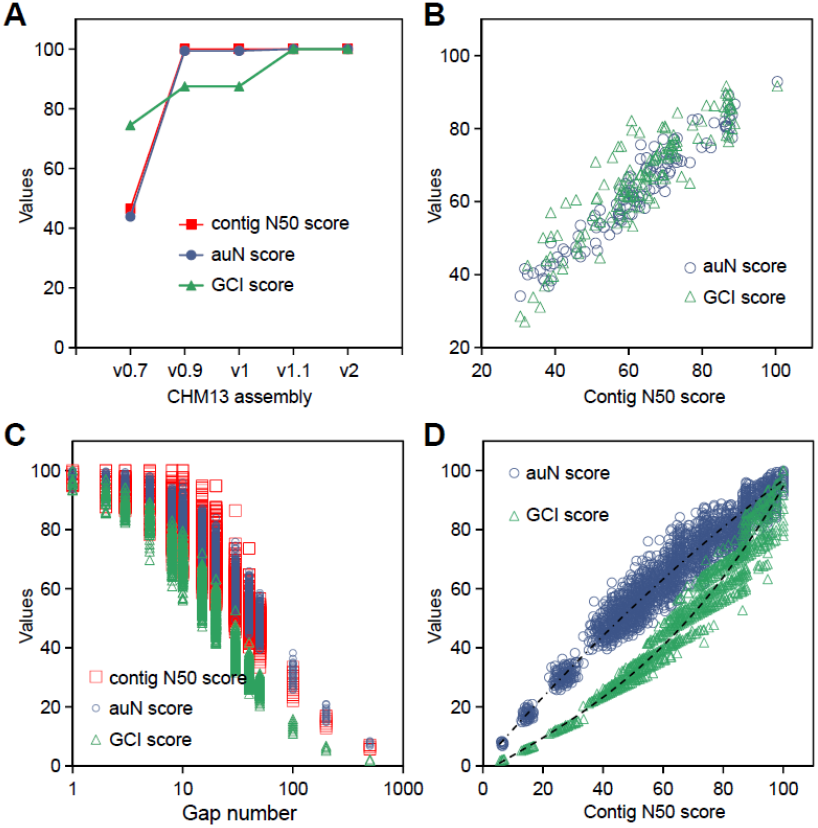
Sensitivity assessment of contig N50, auN and GCI score in quantifying assembly continuity. (A) Contig N50, auN and GCI scores for different versions of CHM13 assembly. Contig N50 and auN were standardized from zero to 100 by their theoretical maximum values, respectively. (B) auN and GCI scores for human phased genome assemblies from HGSVC phase 3. (C) Simulation of different gap numbers (1, 2, 3, 5, 8, 10, 15, 20, 30, 40, 50, 100, 200 and 500) in a human haploid genome. 200 times of simulation for each gap size were performed. (D) Simulated curves of auN and GCI scores with different contig N50 scores.

### GCI evaluation for human genome assemblies

We firstly evaluated GCI performance using three state-of-the-art human genome assemblies (CHM13, CN1 and HG002). CHM13 and CN1 assemblies were gapless, whose contig N50 and E-size/auN values were the theoretical maximum, while HG002 phased assemblies were fragmented with much lower contig N50 and E-size/auN values (Table 1). GCI evaluation based on ONT and Pacbio long reads varied, where GCI scores based on ONT reads were more than twice higher than those using HiFi reads for all three human genomes. This highlights the crucial role of ONT reads in enhancing assembly continuity, in spite of their relatively low base accuracy. Therefore we recommended to use both HiFi and ONT reads in GCI evaluation. By assessing with both HiFi and ONT reads, the curated N50 value of CHM13 assembly that GCI generated reached its theoretical maximum N50 value, while the values were lower than observed contig N50 values calculated from raw assemblies for CN1 and HG002, indicating fewer assembly issues in the CHM13 assembly (Table 1). Consistently, the haploid CHM13 outperformed the other two haplotype-resolved diploid human genomes, with a GCI score of 87.04, compared to CN1 (66.79 for maternal, 77.90 for paternal) and HG002 (18.72 and 27.78). This was expected since less issues were reported in CHM13 assembly due to its high homozygosity.

**Table 1.**
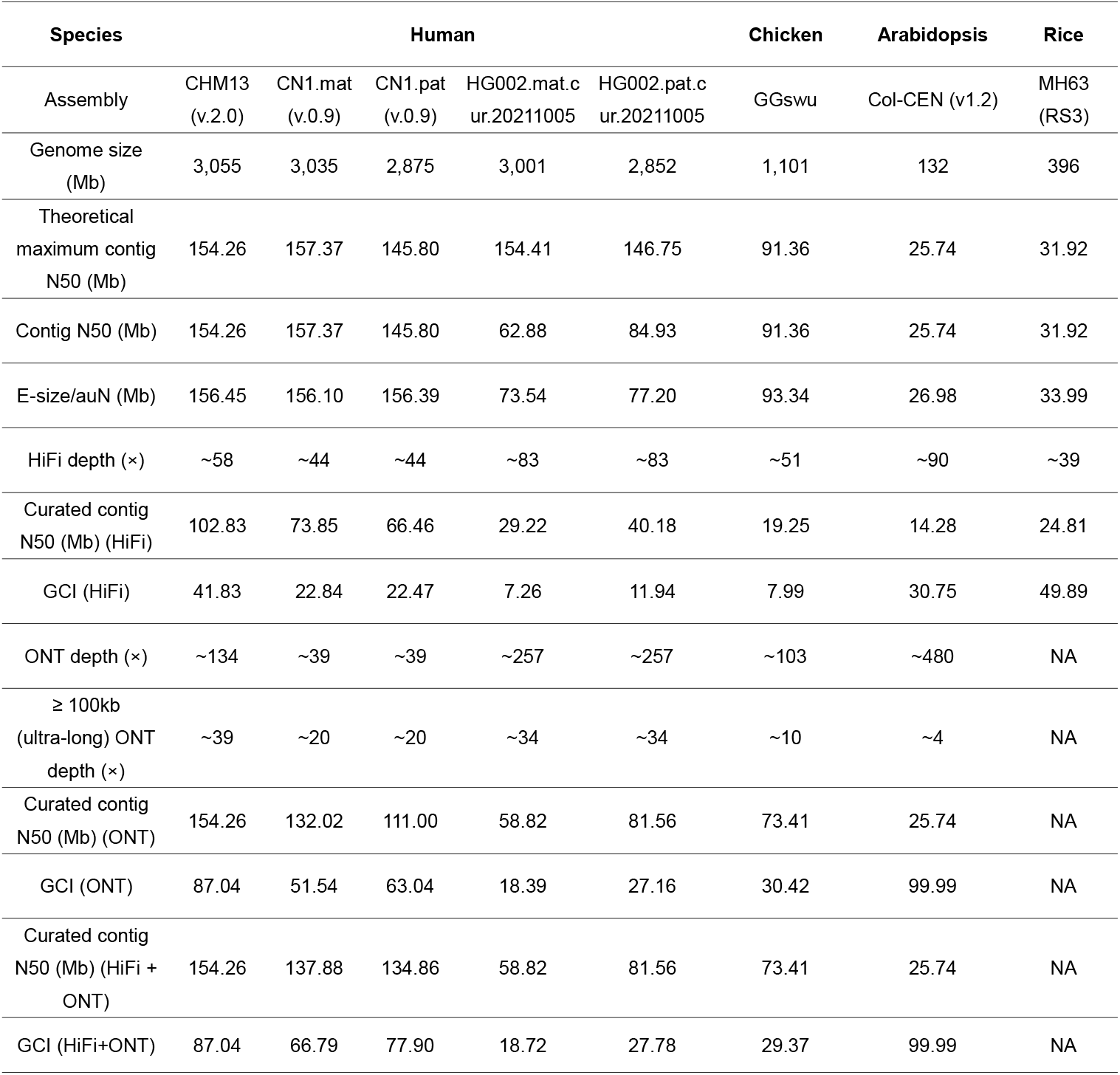
GCI evaluation for the genome assemblies of model species.

Differential GCI sores and high-confidence read mapping supports were observed for two haplotypes in the diploid genomes, which highlights the heterogeneity in genome assembly difficulty due to the potential haplotype-specific complex sequences (Table 1; Fig. S1). While sequencing depth of long reads, especially for ultra-long ONT reads (≥ 100 kb), is crucial for improving the continuity, we noticed that HG002, which was assembled using more HiFi and ONT reads, had lower GCI scores than CN1 (Table 1). This suggests that differential assembly algorithms and gap-filling strategies might contribute to the observed difference in continuity between CN1 and HG002.

Zooming in on the genomic regions of candidate issues in CHM13 assembly, detected by GCI, revealed that all the 11 reported issues were located in the rDNA regions of five acrocentric chromosomes (Chr13, Chr14, Chr15, Chr21 and Chr22)(Figure 3). Notably, no high-confidence read mapping supports were observed in these regions (e.g. the longest issue spanning on Chr13 from ∼6.01 to ∼8.78 Mb). The five rDNA regions are generally recognized as unresolved regions in all the current available human genome assemblies, regardless of whether it is a haploid assembly or diploid phasing assembly (Nurk et al., 2023; Yang et al., 2023). Similarly, for CN1, the identified issues (76 and 63 for maternal and paternal, respectively) were significantly enriched in centromere regions (68, *P* = 8.84e-90 for maternal; 58, *P* = 2.52e-79 for paternal) (Fig. S1), which suggested genomic regions that need to be addressed urgently next by utilizing more advanced sequencing technology and powerful assembling algorithm. In the genome of HG002, a total of 103 and 450 issue regions were identified, including 37 and 364 centromeric issues, for maternal and paternal haplotype, respectively (Fig. S2).

**Figure 3.**
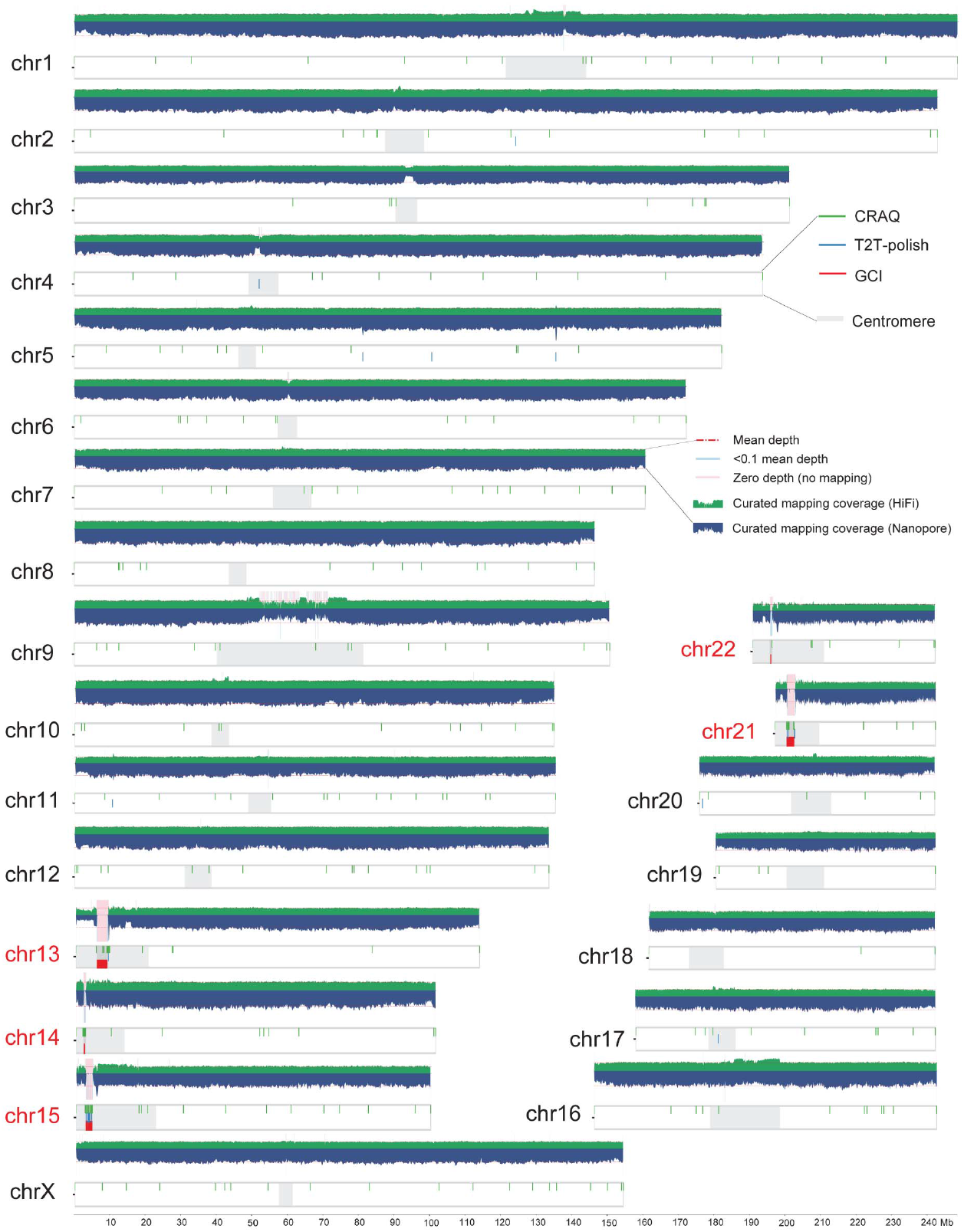
Assembly quality evaluation for human genome CHM13. Long read mapping and assembly issues reported by GCI, T2T-polish, and CRAQ on CHM13 genome. Horizontal red dashed lines in coverage plots represent the whole-genome mean mapping depth. Light blue and pink shadows suggest the regions with low high-confidence read mapping supports (less than 0.1*mean depth) and no support (zero depth), respectively. Chromosomes in red are the five acrocentric chromosomes with rDNA regions.

### GCI evaluation for genome assemblies of non-human model species

*Arabidopsis thaliana* and rice (*Oryza sativa*) are model species of dicot and monocot plants, respectively. For the *Arabidopsis* T2T assembly Col-CEN, its curated N50 value using HiFi reads (14.28 Mb) was significantly lower than that using ONT reads (25.74 Mb), reaching its theoretical maximum. The whole-genome GCI score of Col-CEN reached up to 100 when integrating both HiFi and ONT data (Table 1), which was most likely due to the nature of its compact and simple genome structure (135 Mb with only five chromosomes). The potential gaps in the curated assembly were only detected near the telomeric region of chromosome 2, where a 45S rDNA region was located (Fig. S3). Notably, extremely high coverage was observed in the region from ∼3.3 to ∼3.6 Mb on chromosome 2, due to the presence of mitochondrial insertion, which was flagged as an issue by T2T-polish but not by GCI or CRAQ (Fig. S3).

Rice genome assembly MH63RS3 was the first gapless genome assembled in plants using PacBio reads (Song et al., 2021). Evaluation using its HiFi reads yielded a GCI score of 49.89 (Table 1). The curated N50 value was 80% of its theoretical maximum, which implied the presence of several potential assembly issues. GCI detected a total of 21 issue loci. Within them, issues were absent in centromere and 5S rDNA regions, but observed in the 45S rDNA region on the distal end of chromosome Chr09 (Figure 4A).

**Figure 4.**
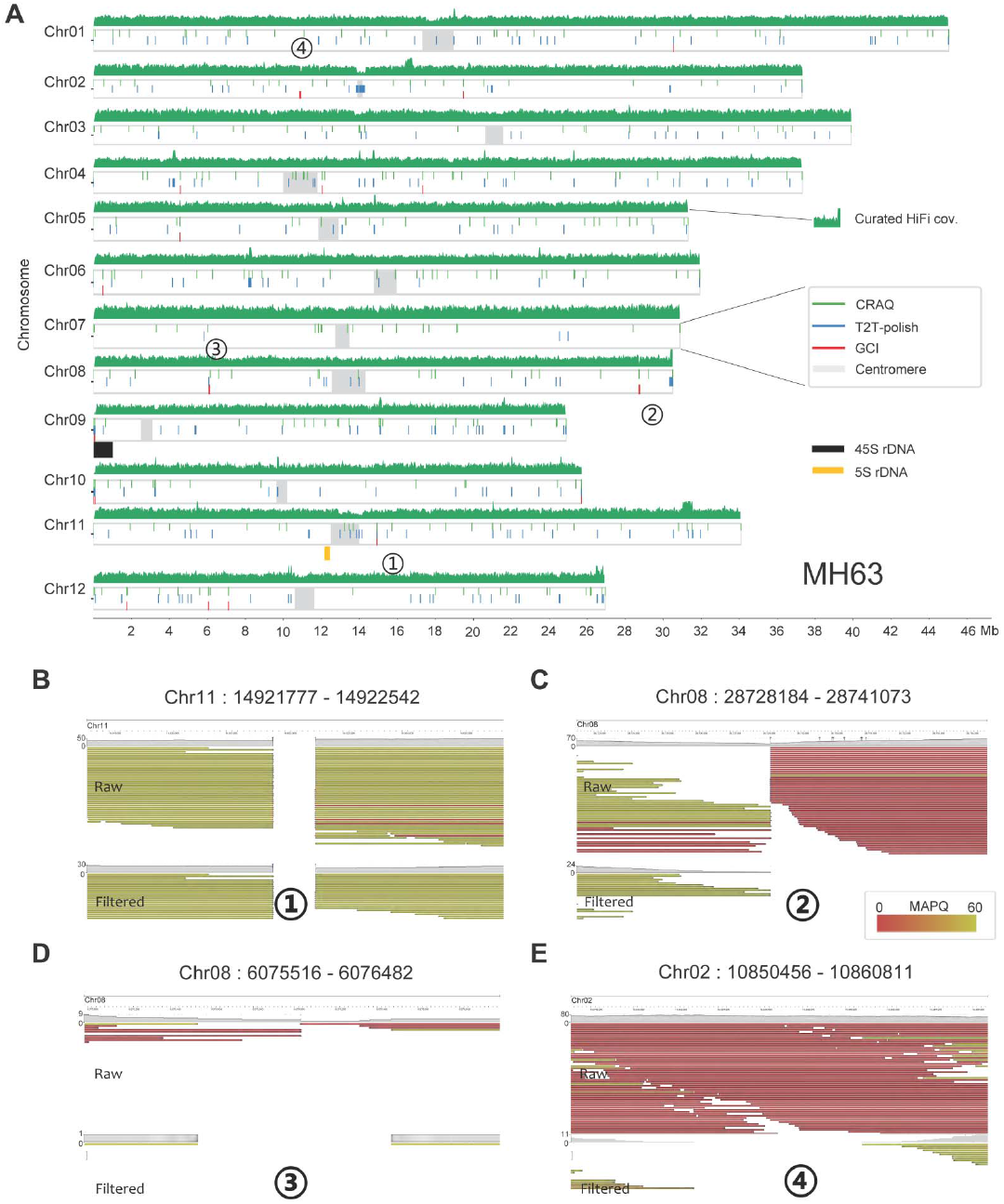
Assembly quality evaluation for rice genome MH63. (A) Genome-wide issues of rice genome assembly MH63RS3 reported by GCI, T2T-polish, and CRAQ. HiFi mapping depth was plotted in sliding 1000 windows on each chromosome. (B) to (E) Genome browser screenshots of four assembly issues in MH63RS3 assembly. ONT reads alignment before and after filtration by GCI were shown.

Unlike the considerable number of so-called T2T genome assemblies in plants, there is relatively limited number of animal genomes have been reported to be completely assembled. For the recently released chicken complete genome assembly (GGswu), we assessed the quality with both HiFi and ONT data and obtained a GCI score of 29.37 and a curated N50 value of 73.41 Mb, which is 80% of its theoretical maximum (Table 1). Totally 582 issues across the whole genome were detected by GCI, spanning a total of 6.94-Mb regions, corresponding with the low GCI score (Fig. S4). In detail, 123 issues (1.12 Mb) were located on 10 macrochromosomes and 19 microchromosomes (totally 1,046 Mb in length), primarily distributed on the ending telomeric regions of chromosomes. Moreover 434 issues (5.59 Mb) were detected on 10 dot chromosomes (40.7 Mb in total length), which implied that further work is needed to validate the assembly accuracy of dot chromosomes.

### Comparison with other tools

We compared the performance of CRAQ and T2T-polish in reporting assembly errors against GCI.

Overall, GCI reported significantly fewer issues than those detected by CRAQ and T2T-polish (Table 2), because the strategy of GCI in detecting assembly issues has avoided a certain number of false positives introduced by non-assembly factors (e.g. sequencing and aligning bias). The CHM13 assembly is a well-recognized complete genome with few issues except for the rDNA regions, which is approved by limited issues reported by GCI (11/11 in rDNA regions) and T2T-polish pipeline (19/27 in rDNA regions). However CRAQ identified up to 328 issues (43 in rDNA regions) (Figure 3), most of which should be false positives. For *Arabidopsis* genome, the 45S rDNA issues on the end of Chromosome 2 were detected by all three tools. The mitochondrial insertion sequences (close to centromere of Chromosome 2) were misidentified as an issue by T2T-polish, owing to its sensitivity in mapping coverage. The chicken assembly exhibited hundreds of assembly issues, using either tool, reflecting its relatively low quality at T2T level (Table 2; Fig. S4).

**Table 2.** Numbers of assembly issues detected by GCI, CRAQ and T2T-polish pipeline for model species genomes.

| Species | Assembly (version) | GCI (No./Length) | CRAQ (No.) | T2T-polish (No./Length) |
| --- | --- | --- | --- | --- |
| Human | CHM13 (v2) | 11 / 5.99 Mb | 328 | 27 / 0.29 Mb |
| Chicken | GGswu | 582 / 6.94 Mb | 1683 | 1829 / 44.67 Mb |
| *Arabidopsis* | Col-CEN (v1.2) | 4 / 19.61 Kb | 25 | 25 / 676.74 Kb |
| Rice | MH63 (RS3) | 21 / 101.77 Kb | 328 | 263 / 1,733.06 Kb |

For the rice genome assembly MH63 (RS3), 21 issues were detected by GCI, among which 15 were overlapped with CRAQ and seven with T2T-polish issues (Table 2). The 45S rDNA region, located on the end of Chr09, showed issue signals by all three tools, while 5S rDNA region on Chr11 reported no issues by either tool (Figure 4A). We manually zoomed in issue regions in genome browser to manually check the assembly quality. Issue Chr11:14,921,777-14,922,542 was one of the five issues shared by all three tools (Figure 4B). No high-confidence read alignment spanned over this region and evident clipping signals were observed, suggesting a gap here.

Issue Chr08:28,728,184-28,741,073 was shared by GCI and CRAQ (Figure 4C). Clipping information was captured by CRAQ to call this issue, and in GCI workflow, removing clipped and low-quality alignments output a gap here. Issue Chr08: 6,075,516-6,076,482 was shared by GCI and T2T-polish (Figure 4D). Limited reads were aligned against this region, which was therefore considered as an issue by T2T-polish pipeline. After filtration by GCI, no reads covered this region, therefore GCI also reported this region as an issue. For GCI-specific issues, most were identified due to the lack of high-confidence read alignment support. For example, in issue region Chr02:10,850,456-10,860,811, no high-confidence read spanned over here, which thus was reported as an issue by GCI (Figure 4E).

## Discussion

Producing a truly complete, contiguous, and accurate genome sequence is the ultimate goal of genome assembly efforts. Wide application of long-read sequencing makes it feasible to obtain a high-quality assemblies, even T2T assembly. The commonly used quality metrics (e.g. N50/NG50/L50/auN, BUSCO/CEGMA, and QV) have been insufficient to distinguish nearly complete genome assemblies whose contig N50 values all reach to this species’ theoretical maximum (i.e. chromosome N50), thus an assembly quality inspector at a higher resolution is required to reveal the potential assembly errors and detect the gaps that would affect the completeness of the assembly. Here we introduced GCI, a genome assembly quality evaluator at single-base resolution, to assess assembly continuity, by integrating long reads. Compared to CRAQ (Li et al., 2023), which collects clipping information of alignments to call assembly errors, GCI uses clipping information to filter alignments. Unlike the Flagger (Liao et al., 2023) and T2T-polish pipeline (Cartney et al., 2022), GCI is not sensitive to reads mapping coverage. In other words, CRAQ and T2T-polish pipeline call the assembly issues by processing abnormal or outlier clipping and depth signals, while GCI collects high-confidence continuous reads that support the correctness of local assembly. Although long-read sequencing is independent of PCR amplification and avoids GC bias, its sequencing bias can still be observed in complex repetitive regions. For example, HSat regions show coverage bias when HiFi (Pacbio Sequel II) and ONT reads are mapped, where mapping coverage decreases to half of whole-genome average depth for HiFi reads but doubled for ONT reads in the DYZ regions (Rhie et al., 2023). Therefore, long-read sequencing bias is a non-negligible factor to introduce false positives of assembly issues detected based on reads coverage. Also the performance of long reads aligning in repetitive regions varies using different aligners (Jain et al., 2022) and unsuitable use of aligning tools might cause the occurrence of mapping anomalies. Based on above considerations, GCI incorporates the alignments from multiple aligners and multiple long-read sequencing platforms. Usually, GCI reports fewer issues than CRAQ and T2T-polish pipeline. However, GCI provides more informative and directional coordinates for subsequent manual check. It should be noted that potential assembly issues reported by GCI, CRAQ or T2T-polish may include misidentification, given that algorithms in genome assembling and reads aligning vary to some extent. In other words, a correct assembly sometimes could be never approved by reads mapping due to the highly repetitive sequence characteristics. For example, high-confidence read alignment supports are usually not observed on 45S rDNA repeat sequences or telomere sequences but this does not mean such assemblies are always incorrect.Therefore a sequence feature-aware assessment method is required to evaluate local assembly quality. The current version of GCI can not capture issues arising from assembly collapse, while CRAQ and T2T-polish can identify such issues through clipping information or mapping depth. Therefore, these quality evaluators are complementary to each other to fully identify all categories of assembly issues, and contribute collectively to the improvement of genome assembly quality together.

## Acknowledgements

The authors thank all members of APG project for helpful suggestions for this work.

## Author contributions

D.W. and G.Z. conceived and designed this study. Q.C. programmed the workflow and implemented the benchmarking and comparison. C.Y. and G.Z. supervised this study. D.W. and Q.C. wrote the original draft. C.Y. and G.Z. revised it. All authors have read and approved the final manuscript.

## Supplementary data

Supplementary data are available online.

## Conflict of interest

None declared.

## Funding

This work is supported by China National Postdoctoral Program for Innovative Talents (BX20220269), and the China Postdoctoral Science Foundation (No. 2023M743045) to D.W.

## Data and code availability

The GCI code is freely available on Github (https://github.com/yeeus/GCI) under the MIT open source license. Detailed issue regions for all assemblies evaluated in this study are available at https://github.com/yeeus/GCI/tree/main/benchmark. For human genome CHM13, its raw reads and different versions of assembly were obtained from Github (https://github.com/marbl/CHM13). All raw reads for HG002 were downloaded from https://github.com/human-pangenomics/HG002_Data_Freeze_v1.0, and the assemblies were downloaded from NCBI under the BioProjects PRJNA794175 and PRJNA794172. For CN1, the updated assembly version v0.9 was obtained from https://genome.zju.edu.cn. ONT and Pacbio long reads for *Arabidopsis thaliana* genome Col-CEN (v1.2) were downloaded from ArrayExpress (accession E-MTAB-10272) and ENA (BioProject PRJEB46164), respectively. Illumina reads of Col accession were downloaded from NGDC (CRA004538) and its assembly was obtained from Github (https://github.com/schatzlab/Col-CEN/tree/main/v1.2). For rice genome MH63, the raw reads and assembly (RS3) were downloaded from NCBI (SRX6957825, SRX6908794, SRX6716809, and SRR13285939) and NGDC (BioProject PRJCA005549), respectively. The assembly and raw reads for chicken genome GGswu are available in NCBI (BioProject accession PRJNA693184). Human genome assemblies from HGSVC phase 3 were downloaded from The International Genome Sample Resource (https://ftp.1000genomes.ebi.ac.uk/vol1/ftp/data_collections/HGSVC3/).

## Supplementary Figures

**Fig. S1.**
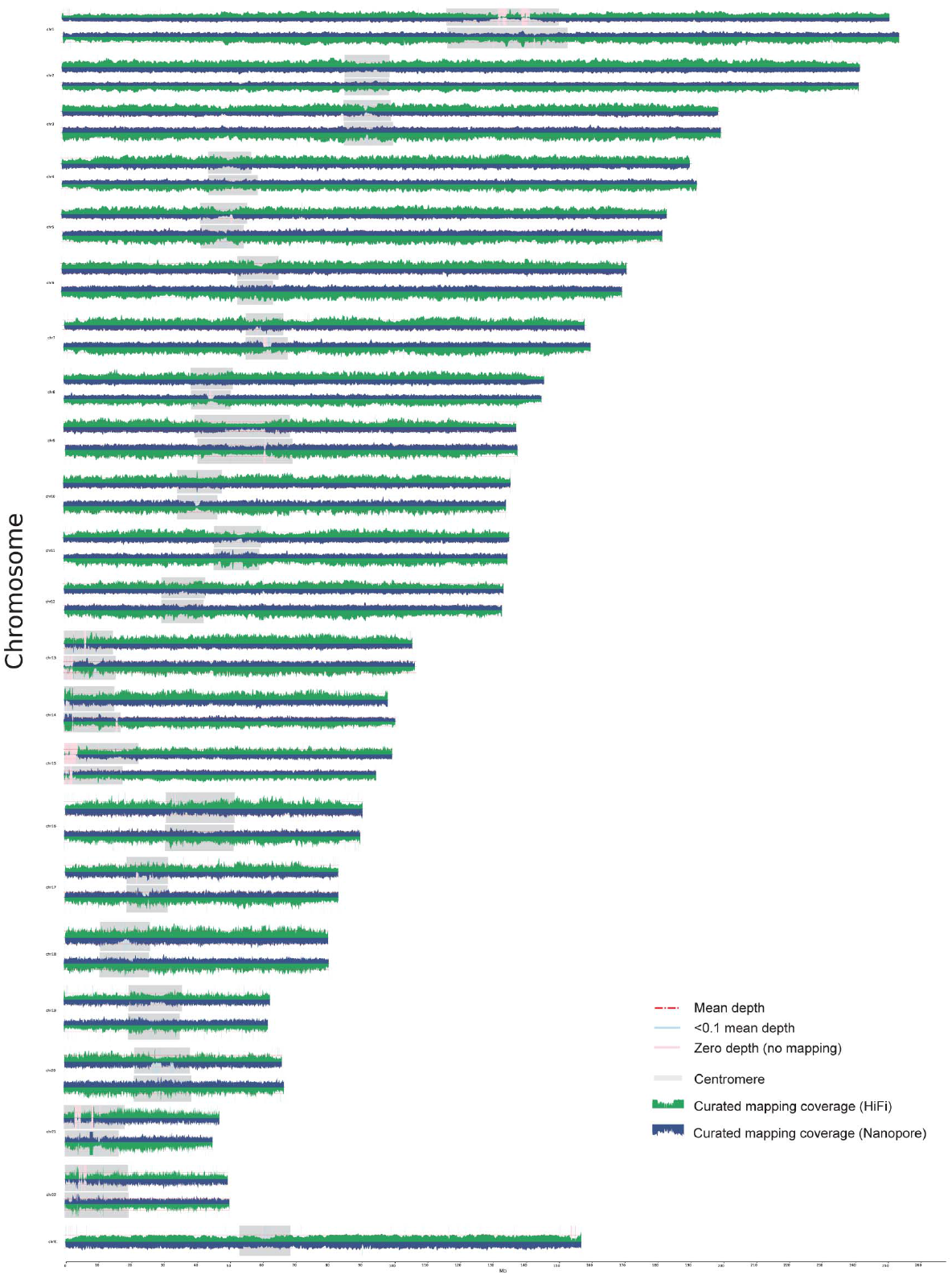
Assembly quality evaluation for human genome CN1 using GCI. Upper and lower tracks for one chromosome represent the mapping coverage of paternal and maternal assembly, where green and blue plots represent the coverage of HiFi and ONT read alignments. Horizontal red dashed lines indicate the whole-genome mean mapping depth. Light blue and pink shadows suggest the regions with low high-confidence read mapping supports (less than 0.1*mean depth) and no support (zero depth), respectively.

**Fig. S2.**
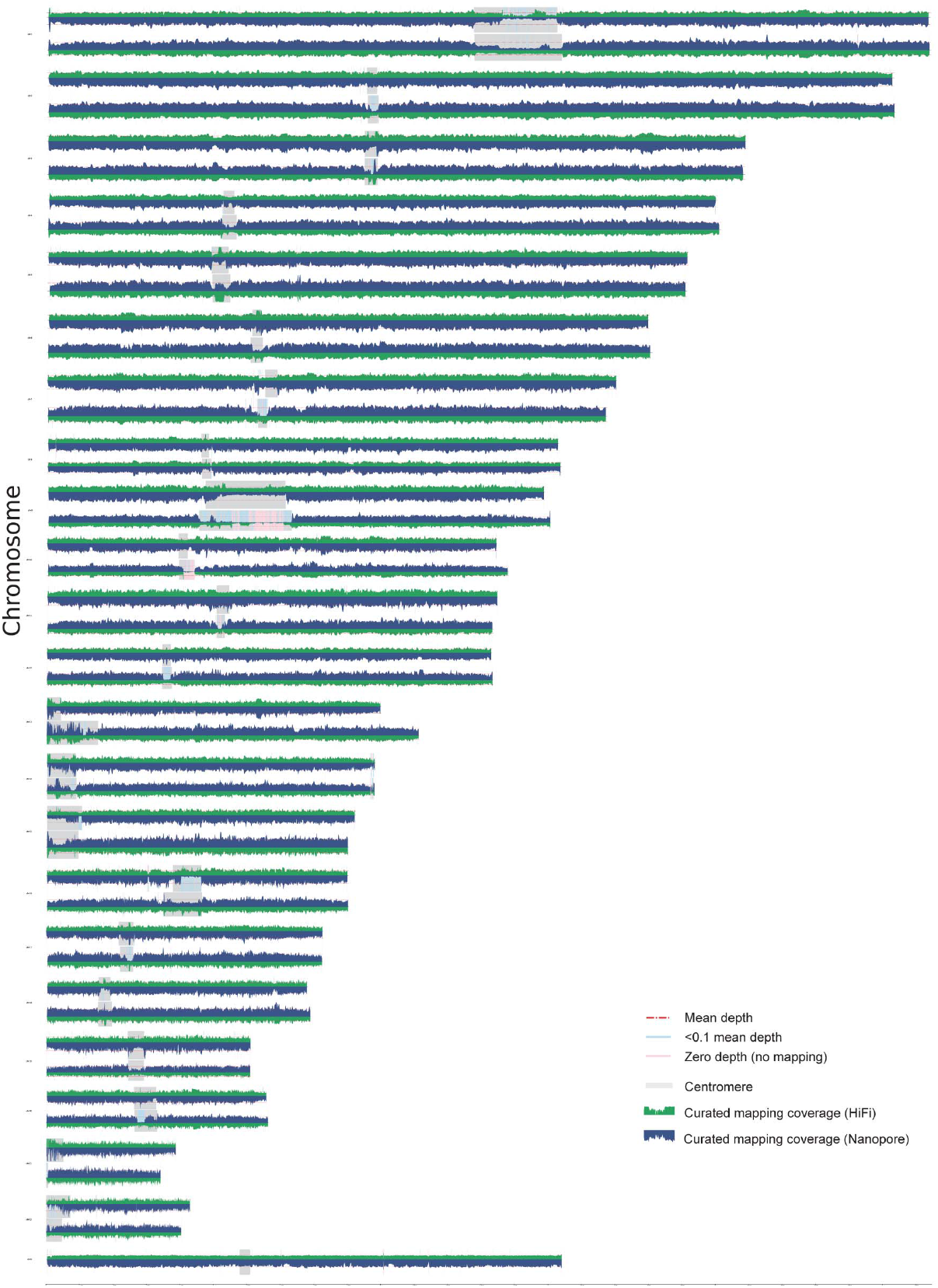
Assembly quality evaluation for human genome HG002 using GCI. Upper and lower tracks for one chromosome represent the mapping coverage of paternal and maternal assembly, where green and blue plots represent the coverage of HiFi and ONT read alignments. Horizontal red dashed lines indicate the whole-genome mean mapping depth. Light blue and pink shadows suggest the regions with low high-confidence read mapping supports (less than 0.1*mean depth) and no support (zero depth), respectively.

**Fig. S3.**
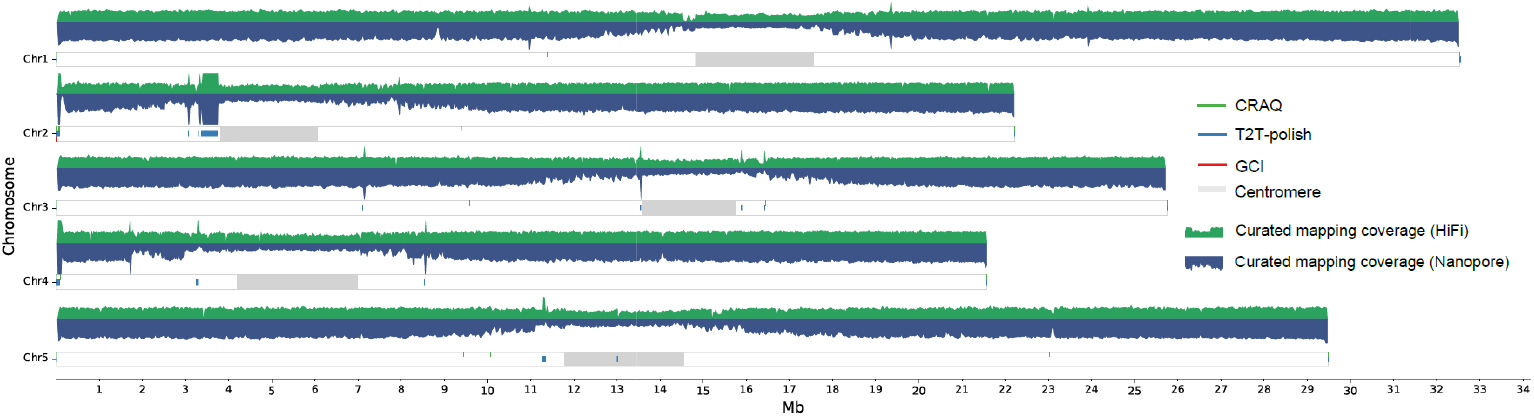
Assembly quality evaluation for *Arabidopsis* genome Col-CEN using GCI, T2T-polish and CRAQ.

**Fig. S4.**
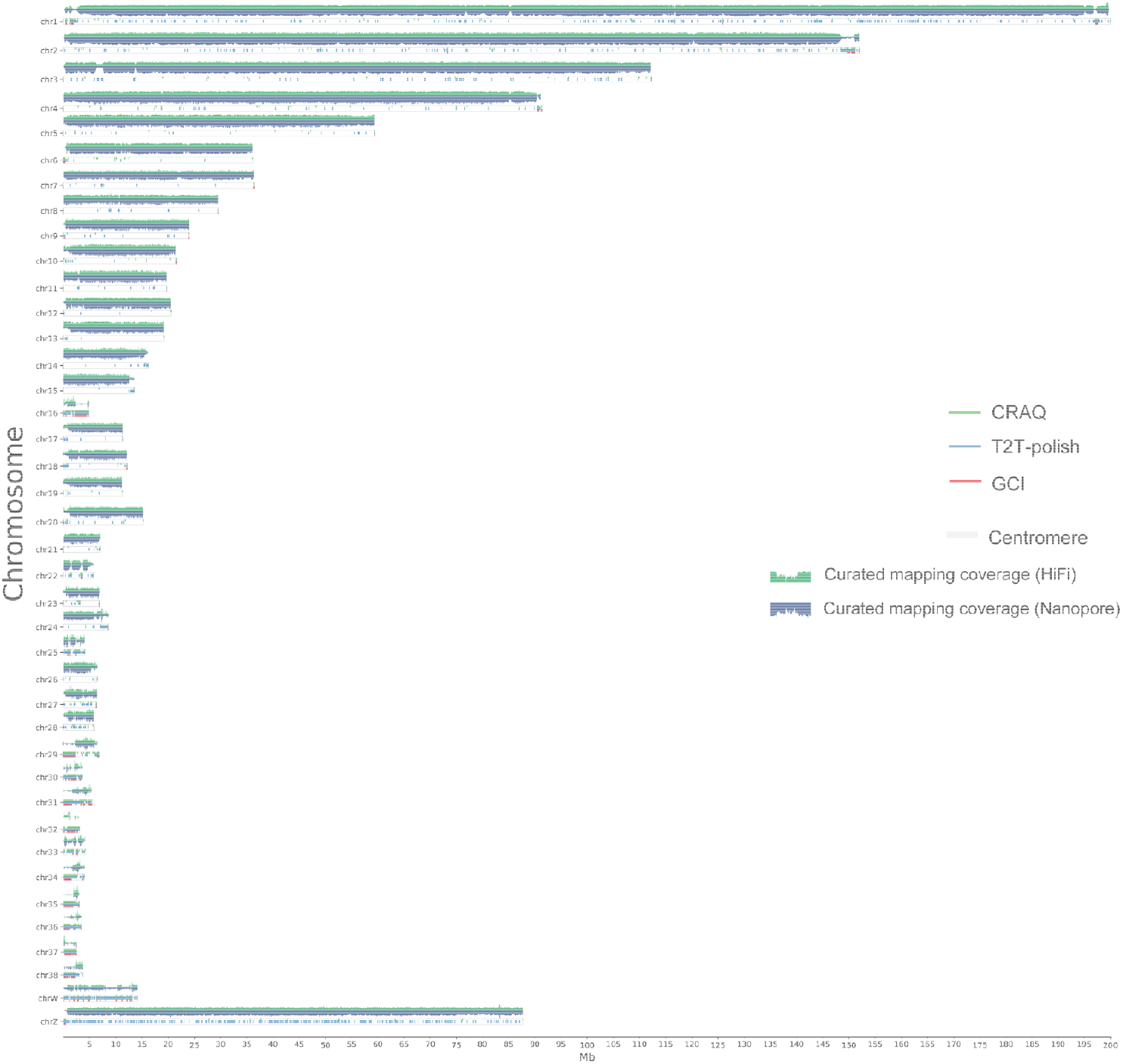
Assembly quality evaluation for chicken genome GGswu using GCI, T2T-polish and CRAQ.

## Notes

https://github.com/yeeus/GCI

### Competing Interest Statement

The authors have declared no competing interest.

## References

Bzikadze, A.V. et al.(2022) Fast and accurate mapping of long reads to complete genome assemblies with VerityMap.Genome Res., 32, 2107–2118.

Huang, Z. et al.(2023) Evolutionary analysis of a complete chicken genome.Proc. Natl. Acad. Sci., 120, e2216641120.

Jain, C. et al.(2022) Long-read mapping to repetitive reference sequences using Winnowmap2.Nat. Methods, 19, 705–710.

Jarvis, E.D. et al.(2022) Semi-automated assembly of high-quality diploid human reference genomes.Nature, 611, 519–531.

Li, H. (2018) Minimap2: pairwise alignment for nucleotide sequences.Bioinformatics, 34, 3094–3100.

Li, K. et al.(2023) Identification of errors in draft genome assemblies at single-nucleotide resolution for quality assessment and improvement.Nat. Commun., 14, 1–12.

Liao, W.-W. et al.(2023) A draft human pangenome reference.Nature, 617, 312–324.

Mc Cartney, A.M. et al.(2022) Chasing perfection: validation and polishing strategies for telomere-to-telomere genome assemblies.Nat. Methods, 19, 687–695.

Naish, M. et al.(2021) The genetic and epigenetic landscape of the Arabidopsis centromeres.Science,374, eabi7489.

Nurk, S. et al.(2020) HiCanu: accurate assembly of segmental duplications, satellites, and allelic variants from high-fidelity long reads.Genome Res., 30, 1291–1305.

Nurk, S. et al.(2022) The complete sequence of a human genome.Science, 376, 44–53.

Rhie, A. et al.(2020) Merqury: reference-free quality, completeness, and phasing assessment for genome assemblies.Genome Biol., 21, 245.

Rhie, A. et al.(2023) The complete sequence of a human Y chromosome. Nature, 621, 344–354.

Salzberg, S.L. et al.(2012) GAGE: A critical evaluation of genome assemblies and assembly algorithms.Genome Res., 22, 557–567.

Sim, S.B. et al.(2022) HiFiAdapterFilt, a memory efficient read processing pipeline, prevents occurrence of adapter sequence in PacBio HiFi reads and their negative impacts on genome assembly.BMC Genomics, 23, 157.

Song, J.-M. et al.(2021) Two gap-free reference genomes and a global view of the centromere architecture in rice.Mol. Plant, 14, 1757–1767.

Wang, P. and Wang, F. (2023) A proposed metric set for evaluation of genome assembly quality.Trends Genet., 39, 175–186.

Yang, C. et al.(2023) The complete and fully-phased diploid genome of a male Han Chinese.Cell Res., 1–17.

